# Heritability of the structures and ^13^C fractionation in tomato leaf wax alkanes: a genetic model system to inform paleoenvironmental reconstructions

**DOI:** 10.1101/110718

**Authors:** Amanda L.D. Bender, Daniel H. Chitwood, Alexander S. Bradley

## Abstract

Leaf wax *n*-alkanes are broadly used to reconstruct paleoenvironmental information. However, the utility of the *n*-alkane paleoclimate proxy is modulated by the extent to which genetic as well as environmental factors influence the structural and isotopic variability of leaf waxes. In paleoclimate applications, there is an implicit assumption that most variation of leaf wax traits through a time series can be attributed to environmental change and that biological sources of variability within plant communities are small. For example, changes in hydrology affect the δ^2^ H of waxes though rainwater and the δ^13^C of leaf waxes by changing plant communities (i.e., C_3_ versus C_4_ input). Here we test the assumption of little genetic control over 5 C variation of leaf wax by presenting the results of an experimental greenhouse growth study in which we estimate the role of genetic variability on structural and isotopic leaf wax traits in a set of 76 introgression lines (ILs) between two interfertile *Solanum* (tomato) species: *S. lycopersicum* cv M82 (hereafter cv M82) and *S. pennellii.* We found that the leaves of *S. pennellii,* a wild desert tomato relative, produces significantly more *iso*-alkanes than cv M82, a domesticated tomato cultivar adapted to water-replete conditions; we introduce a methylation index to summarize the ratio of branched (*iso*- and *anteiso-*) to total alkanes. Between *S. pennellii* and cv M82, the *iso*-alkanes were found to be enriched in ^13^C by 1.2–1.4%o over *n*-alkanes. By modeling our results from the ILs, we report the broad-sense heritability values (*H*^2^) of leaf wax traits to describe the degree to which genetic variation contributes to variation of these traits. Individual carbon isotope values of alkanes are of low heritability (*H*^2^ = 0.13–0.19), suggesting that δ^13^C of leaf waxes from this study are strongly influenced by environmental variance, which supports the interpretation that variation in the 5 C of wax compounds recorded in sediments reflects paleohydrological changes. Average chain length (ACL) values of *n*-alkanes are of intermediate heritability (*H*^2^ = 0.30), suggesting that ACL values are strongly influenced by genetic cues.

## 1 Introduction

Long chain (C_21_ – C_39_) *n*-alkanes are characteristic components of the cuticular waxes of terrestrial plants (Jetter et al., 2006). Alkanes are geologically stable, and their structures and isotopic compositions carry biological and environmental information. In a geological context, this information can be used for paleoenvironmental and paleoecological reconstructions. Structural traits in w-alkanes, such as average chain length (ACL), may relate to climatic variables such as temperature and humidity, as well as to the plant sources of the *n*-alkanes (Bush and McInerney, 2013, 2015). Stable isotopes of carbon (δ^13^C) in plant materials, including waxes, relate to the plant's carbon fixation pathway (Naafs et al., 2012; Tipple and Pagani, 2010), to physiological parameters of plants such as water use efficiency (WUE) and stomatal conductance (Easlon et al., 2014), and to environmental parameters such as atmospheric CO_2_ concentration (Schubert and Jahren, 2012). Hydrogen ratios (δ^2^H) in plant wax *n*-alkanes relate to the δ^2^H of rainwater, as well as to a number of environmental and physiological parameters (Sachse et al., 2012).

Although these characteristics are informative, the utility of *n*-alkanes in tracing environmental variability is moderated by uncertainty about the degree to which structural and isotopic variability is a function of plant biology in addition to environmental conditions. Therefore, the efficacy of sedimentary *n*-alkanes in paleoclimate applications can be informed by a better understanding of plant biology, and of the genetic and physiological factors that control structural and isotopic variations in plant wax compounds.

One approach to understanding the role of biological variability consists of sampling leaf wax material from a range of species in a terrestrial environment. For example, Hou et al. (2007a) showed that δ^2^H values co-vary with δ^13^C values of leaf waxes and may be related to WUE in a range of tree species sampled in a single environment near Blood Pond, Massachusetts (Hou et al., 2007a). Leaf wax δ^2^H values of different plant types from the same environment have been shown to vary by as much as 70%o (Hou et al., 2007b), and interspecies variations with a standard deviation of 21%o were observed in the hydrogen isotopic composition of plant-derived w-alkanes in an arid ecosystem (Feakins and Sessions, 2010). Total *n*-alkane abundances vary greatly among angiosperms; for example, total *n*-alkane abundances in different species of the same plant family (Betulaceae) range from <50 μg/g dry leaf to 1300 μg/g dry leaf (Diefendorf et al., 2011). Carbon isotope values of *n*-alkanes measured from a wide range of angiosperms have been reported to vary by up to ~10‰ (Diefendorf et al., 2011). These examples demonstrate that biological variability is present among various lineages. In some cases this is strongly expressed. For example, differences in photosynthetic pathways impart a strong carbon isotopic discrimination in *n*-alkanes. Across 10 studies, *n*-C_29_ and *n*- C_31_ alkanes from C3 plants showed mean δ^13^C values of −34.0% and −34.3% with 1σ variation of 3.3% and 3.0%, respectively, whereas the same alkanes from C4 plants have mean δ^13^C values of −21.4% and −21.7% and 1σ variations of 2.3% and 2.3% in C4 plants (Figure 1; Supplementary Dataset 1). Within families within the broader class of C3 plants, biological variability might be expected to have an effect on δ^13^C values of alkanes. This could in principle be true of variability even within a single plant genus.

**Figure 1:**
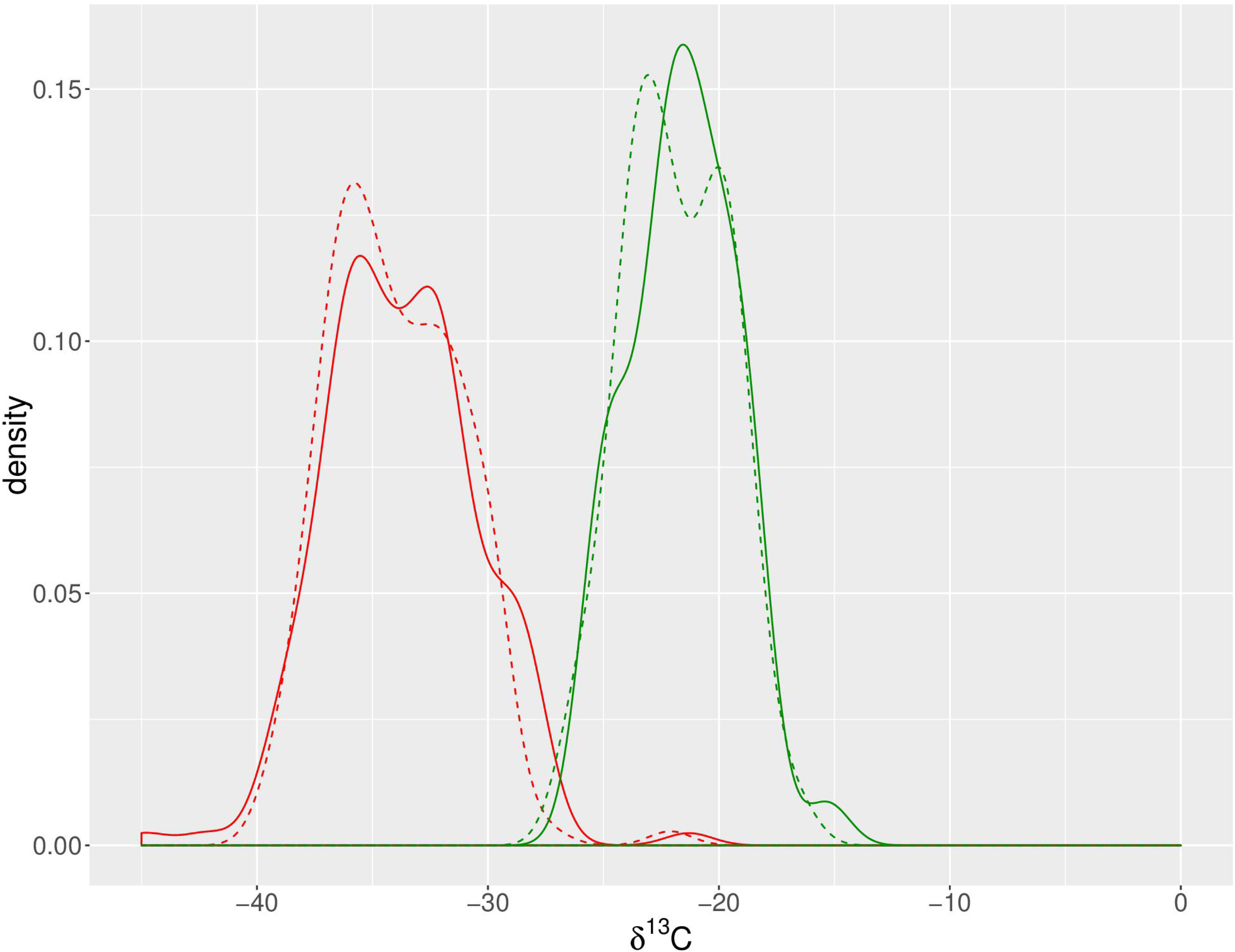
Density plot of *n*-C_29_ and *n*-C_31_ alkane δ^13^C values from C_3_ and C_4_ plants. *n*-C_29_ (solid lines) and *n*-C_31_ (dashed lines) alkanes from C3 plants (red) show a 1σ variation of 3.3‰ and 3.0 ‰ respectively, while the same alkanes have 1σ variations of 2.3‰ and 2.3‰ in C4 plants (green). Data shown are from 10 published studies (see Supplementary Dataset 1).

Studies of biological variability of *n*-alkane traits provide an estimate for the magnitude of potential variation but do not provide information regarding the mechanistic processes underlying that variation. An enhanced understanding of the mechanisms responsible for isotopic variability of *n*-alkanes could allow for more precise reconstructions of precipitation and/or temperature. Mechanistic questions might be addressed by studies that examine the variation of leaf wax traits with respect to physiology (e.g., Gao et al., 2015; Smith and Freeman, 2006; Tipple et al., 2012) or genetics (e.g., Gao et al., 2014).

Genetic approaches are particularly relevant for honing the understanding of leaf wax trait variability. Continuous phenotypic traits, such as the isotopic composition of *n*-alkanes, can be parameterized as reflecting a combination of genotypic factors that interact with the environment. Although this interaction is commonly recognized in biological studies, the genotype-environment interaction is consistently neglected in paleoenvironmental applications. In paleoenvironmental reconstructions that employ hydrogen isotopes of leaf wax compounds, isotopic variation is implicitly assumed to be mostly or entirely a function of environmental variability. Similarly, variation in the carbon isotopic composition of leaf waxes is usually attributed to differing inputs of C3 and C4 plants (Castañeda et al., 2009a, 2009b; Eglinton et al., 2002; Feakins et al., 2005, 2007; Tipple and Pagani, 2010). For individual plant species, however, there is also a strong correlation between the ^13^C content of leaf waxes and mean annual precipitation (Diefendorf et al., 2010). The degree to which genetic variation among and within species contributes to isotopic variation is not well constrained.

This variation can be described using the broad-sense heritability of a trait, a widely used statistic in quantitative genetic studies. Broad-sense heritability (*H*^2^) is defined as the proportion of total phenotypic variance that can be attributed to genetic variation (Futuyma, 1998), as given by the equation:

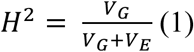

where *V_G_* is genetic variance and *V_E_* is environmental variance. The use of phenotypic traits such as wax δ^13^C or δD values in paleoenvironmental reconstruction implicitly makes one of two assumptions: i) that variation through a time series can be attributed to environmental change, and that biological sources of variability are small, or ii) that differences in e.g. δ^13^C can be modeled as simple mixing between two end members (such as C_3_ and C_4_ plants). In this study we assess the validity of the first assumption.

We directly assess the broad-sense heritability of structural and isotopic leaf wax traits in a greenhouse using a model species complex consisting of precisely defined near-isogenic introgression lines (ILs; Eshed and Zamir, 1995) between two interfertile *Solanum* (tomato) species. Each of the 76 ILs possess a single introgressed genomic segment from the desert wild tomato relative *Solanum pennellii* in a domesticated tomato *Solanum lycopersicum* cv M82 background (hereafter, cv M82). Together, the introgression segments of the 76 ILs span the entire domesticated tomato genome. These two species are adapted to regions with vastly different hydrological settings. Endemic to the dry slopes of the Central Peruvian Andes (Warnock, 1991), *S. pennellii* bears smaller fruit and leaves that are smaller and less complex than those of cv M82, which was selected during cultivation in water-replete conditions (Chitwood et al., 2013). The leaves of these two species also vary in their wax content; epicuticular lipids comprised 0.96% and 19.9% of total leaf dry weight in 17-week old leaves from cv M82 and *S. pennellii,* respectively (Fobes et al., 1985). The genome of *S. pennellii* has been well studied and sequenced (Bolger et al., 2014), which improves the utility of this model organism for genetic study.

Although these species are not abundant producers of leaf waxes in terrestrial ecosystems, they nonetheless provide a useful tool for investigating plant genetics and physiology. These can be considered model organisms in the same way that *Escherichia coli* is used as a model organism for understanding fundamentals of bacterial physiology and genetics. We use this model species complex to determine the role of heritability in the production of plant wax traits that are central to paleoclimatic reconstruction. This approach allows us to test the implicit assumption that genetic variance plays a limited role in driving variation of leaf wax traits that are preserved in sediments through time, and whether the recorded variation may reflect a high-fidelity paleoclimate signal.

In this study, we use the *Solanum pennellii* ILs to resolve genetic and environmental effects on leaf wax δ^13^C values and structural traits. *n*-Alkanes are among the most abundant and simplest of waxes to extract and isolate, and are thus commonly analyzed from sediments and modern plants. Among tomatoes and other plants in the *Solanum* genus, the most abundant alkanes are *n*-alkanes, but branched *iso*- and *anteiso*-alkanes have also been identified within the *Solanum* genus (Girard et al., 2012; Silva et al., 2012; Smirnova et al., 2013; Smith et al., 1996; Szafranek and Synak, 2006). Other plants of the Solanaceae family (Grice et al., 2008; Heemann et al., 1983; Rogge and Hildemann, 1994) also contain branched alkanes, as well as members of the Lamiaceae family (Huang et al., 2011; Reddy et al., 2000), Aeonium genus (Eglinton et al., 1962), and an Arctic chickweed (Pautler et al., 2014). Long chain *iso-* and *anteiso-* alkanes are expected to derive from the same biosynthetic pathway as *n*-alkanes, albeit with different biosynthetic precursors that might contribute to systematically distinctive δ^13^C values for these alkane types (Grice et al., 2008).

Although leaf wax traits have been shown to adapt dynamically to environmental stresses (Grice et al., 2008; Kosma et al., 2009), here we establish the static features of leaf wax traits of *S. pennellii,* cv M82, and the *S. pennellii* IL population by growing all plants in the same greenhouse conditions. We identify quantitative trait loci (QTLs) that underlie many leaf wax traits and calculate broad-sense heritability values to estimate the proportion of phenotypic variance attributable to genetic variance. Our results have important implications for uncovering the degree to which we can expect environmental versus genetic factors to modulate variability in leaf wax traits.

## 2 Materials and methods

### 2.1 Plant materials, growth conditions, and experimental design

We obtained second-generation *Solanum pennellii* ILs (Eshed and Zamir, 1995), *Solanum pennellii,* and *Solanum lycopersicum* cv M82 seeds from the Tomato Genetics Resource Center and the lab of Neelima Sinha (University of California, Davis) and Dani Zamir (Hebrew University, Rehovot, Israel). All seeds were prepared and germinated at the Donald Danforth Plant Science Center in St. Louis, MO as described in Chitwood et al. (2013).

#### 2.1.1 Growth conditions for cv M82 and *S. pennellii*

In order to characterize the variance of leaf wax traits between the two parent lines, we grew ten replicates of each parent species in the greenhouse from November 2013 – January 2014. The seeds were germinated in late November 2013. Seedlings were transplanted into 2-gallon planters in the greenhouse and staggered along a greenhouse bench. Seedlings were vigorously top watered after transplanting and further watered and fertilized to ensure plant growth; irrigation water was supplied from a tap water reservoir.

Anthesis began in late December 2013; leaves were collected from each plant in early January 2014. One leaf was collected from each plant for leaf wax extraction and analysis based on specific criteria: (i) the leaf was fully developed (i.e., leaflets were fully unfurled), and (ii) the leaf was young (i.e., close to the top of the plant, arising after the reproductive transition). Each sample comprised the five primary leaflets of each leaf (terminal, distal, and proximal lateral left and right).

#### 2.1.2 Growth conditions for ILs and cv M82

We grew the 76 ILs from December 2013 – February 2014. Seeds were germinated in December and transplanted to 3-gallon planters in the greenhouse in January. The ILs and cv M82 were arranged in a randomized block design with four replicates (Supplemental Figure 1, Supplemental Dataset 2). Watering and fertilization proceeded as with the parent lines.

After anthesis in early February, we collected leaf samples in late February according to identical criteria as with the parent lines, collecting leaves that were 6-8” in length. Our growing efforts were successful for all but four ILs (ILs 2.2, 5.3, 7.1, and 8.2).

During the growth period for the ILs, we monitored ambient conditions of the greenhouse: relative humidity, temperature, *p*CO_2_, and δ^13^C_CO2_. Daytime temperature and relative humidity were monitored with custom systems integrated with the greenhouse (Argus Control Systems, Ltd.). Temperatures were maintained at approximately 78°F (25.6°C). We set up a Picarro Cavity Ring-Down Spectrometer G2131-*i* Analyzer in the greenhouse to monitor *p*CO_2_ and δ^13^C_CO2_; these data were aggregated from 5-minute interval measurements (Supplemental Figure 2, Supplemental Dataset 3).

### 2.2 Leaf harvest and lipid extraction

Each sample consisted of five leaflets (terminal, distal lateral left and right, proximal lateral left and right) from a single leaf of each plant. We measured leaf area from all sample leaflets with a flatbed scanner and ImageJ software (Abràmoff et al., 2004). The collected leaf samples were cut into 1 cm^2^ pieces, placed into pre-baked 15 mL clear borosilicate vials (Qorpak), and then dried in a 70°C oven for 48 hours. We extracted epicuticular waxes from the dried leaf samples by adding 5 mL of hexane (Omni-Solv HR-GC Hexanes 98.5%, VWR International, LLC) and agitating via pumping with a Pasteur pipette. The resulting total lipid extract (TLE) was collected into a separate 15 mL borosilicate vial; the extraction step was repeated three times, and the three extractions were pooled. We evaporated the TLE to dryness with heat (30°C) under a steady stream of nitrogen gas (FlexiVap Work Station, Glas-Col).

To isolate *n*-alkanes for analysis, we performed silica gel column chromatography on the dried TLE of each sample. We transferred the TLE with 50 μ L of hexane to a silica gel column (5 cm × 4 mm Pasteur pipette packed at the base of the taper with a small amount of laboratory grade glass wool that had been previously baked at 550°C for 8 hours, a thin layer of chromatography grade sand [50-70 mesh particle size, baked at 850°C for 8 hours], and filled 3-4 cm high with H_2_O-deactivated silica gel [230-400 mesh particle size, baked at 550°C for 8 hours]). We collected the *n*-alkane fraction by eluting with hexane. The polar compounds retained on the silica gel column were eluted with ethyl acetate and archived.

### 2.3 Leaf wax structural analysis

We analyzed the *n*-alkane fractions using an Agilent 7890A Gas Chomatograph (GC) equipped with a 5975C Series Mass Spectrometric Detector (MSD) system at the Biogeochemistry Laboratory at Washington University in St. Louis. The GC was equipped with an Agilent J&W HP-5ms column (30 m long, 0.25 mm inner diameter, 0.25 μm film thickness). The GC-MSD system was equipped with an Agilent 7650A Automatic Liquid Sampler. The GC oven had an initial temperature of 60°C and was heated at a rate of 6°C/min to the final temperature of 320°C, which was held for 20 minutes. One sample run lasted approximately 65 minutes. *n*-Alkanes were identified by their mass spectra and quantified against an internal standard (n-hexadecane-d_34_, 98 atom%, Sigma-Aldrich).

### 2.4 Carbon isotope analysis

Carbon isotopic compositions of w-alkanes were determined on a gas chromatograph coupled via a combustion reactor to a Thermo-Finnigan Delta Plus mass spectrometer at the Biogeochemistry Laboratory at Washington University in St. Louis. δ ^13^C values were measured against an internal *n*-alkane standard (C_18_) and reported in ‰ against the standard Vienna Pee Dee Belemnite (V-PDB). All samples were analyzed in triplicate. An *n*-alkane standard (B3 or A5) of 15 externally calibrated *n*-alkanes and a fatty acid standard (F8) of 8 externally calibrated fatty acids provided by A. Schimmelmann (Indiana University) were measured between every fifth sample injection. Analytical uncertainty of reported δ^13^C values ranges between ± 0.2% and 0.3% (SEM), dependent on the number of analytical replicates, after propagating the uncertainty of replicate analyses and external molecular standards (Polissar and D'Andrea, 2014) (data available on GitHub repository at http://github.com/aldbender/13C-heritability).

Ambient greenhouse CO_2_ was monitored during growth of the *Solawum* ILs from December 19, 2013 – February 24, 2014, except for the period from January 1-14 when a technical error prevented data collection. For the ILs and cv M82 plants grown simultaneously in the greenhouse, we report the apparent fractionation (^13^ε) between the carbon isotope value of atmospheric CO_2_ (δ^13^C_atm_) and the carbon isotope of the lipid (δ ^13^C_lipid_):

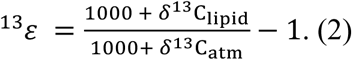

Carbon isotope values of lipids from *S. pewwellii* and cv M82 grown during November 2013 are not reported as Δ values because the carbon isotopic value of ambient greenhouse CO_2_ was not recorded during their growth period. We also report the differences in δ^13^C (‰) between *n*-alkanes and *iso*-alkanes of the same carbon-numbered alkanes, expressed as δ_*n*-alkanes_ – δ_*iso*-alkanes_ (or simply δ_*n*_ – δ_*iso*_).

### 2.5 Characterizing *n*-alkane distributions

We characterized the distribution of alkanes for each sample by calculating a suite of summary traits: the methylation index (a novel measure of this study), and average chain length (ACL) and carbon preference index (CPI) each calculated individually for *n*-, *iso-,* and *anteiso*-alkanes. Here, we define the methylation index as the relative abundance of branched (*iso*- and *anteiso-*) alkanes to the total of branched and unbranched (*normal*) alkanes as in the equation:

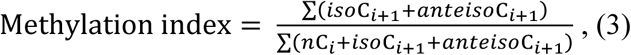

where *iso*C_*i*+1_, *awteiso*C_*i+*1_, and *n*C_*i*_ are the concentrations of *iso-, awteiso-,* and *n-* alkanes with *i* carbon chain length, respectively. A methylation index value of 0 indicates that there are only *n*-alkanes in a sample, whereas a methylation index value of 1 indicates that there are only *iso-* and *awteiso*-alkanes in a sample. The average chain length (ACL) is the weighted average of the carbon chain lengths, defined as:

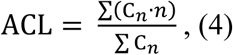

where *C_n_* is the concentration of each alkane with *n* carbon atoms. The carbon preference index (CPI) measures the relative abundance of odd over even carbon chain lengths, where:

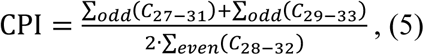

and summarizes the dominance of odd carbon number alkanes over even carbon number alkanes. Greater CPI values indicate a greater predominance of odd over even chain lengths.

### 2.6 Statistical modeling and QTL analysis

Data from all traits measured from the ILs are reported in Supplemental Dataset 3. The R code and data sets used for modeling are available on GitHub at http://github.com/aldbender/13C-heritability. Leaf wax traits were modeled using mixed-effect linear models with the lme4 packages (http://CRAN.R-project.org/package=lme4) in R (R Development Core Team, 2015). Before modeling, we compared the measured values against theoretical normally distributed values in a Q-Q plot to check whether the measured values came from a normally distributed population. If a trait deviated from a normal distribution, we transformed the trait by either taking the square root, log, reciprocal, or arcsine of the trait and tested for normality of each transformed population via the Shapiro-Wilk test, using the transformation that resulted in the least deviation from a normal distribution (see Supplemental Dataset 4 for measured trait summaries and the selected transformation for each trait). In order to perform linear modeling, all δ_*n*_ − δ_*iso*_ values were additionally transformed by calculating the absolute value. After performing the mixed-effect linear modeling, the normal distribution of residuals in the model was verified. We extracted *p*-values for significant (*p* < 0.05) differences between ILs and cv M82 from the models using the pvals.fnc function from the language R package (http://CRAN.R-project.org/package=languageR); these *p*-values were used to generate the QTL analysis. We calculated the broad sense heritability values (*H*^2^) from the estimates of genetic and environmental and residual variances from the mixed-effect linear modeling (see Supplemental Dataset 5), as defined in Equation 1.

### 2.7 Hierarchical clustering of traits

Hierarchical clustering is used to build a hierarchy of traits that cluster together based on dissimilarities between sets of trait observations. We performed a correlation analysis of leaf wax traits from this study with traits existing in the phenomics database (Phenom-Networks, www.phenome-networks.com). For each trait in a data set, data were z-score normalized in order to transform all data ranges to a standardized average and standard deviation. Z-scores were averaged across replicates. We performed hierarchical clustering using the hclust function from the stats package in R (R Development Core Team, 2015), clustering by the absolute value of the Pearson correlation coefficient using Ward's minimum variance method. We created a correlation matrix (Spearman) between all traits measured in this study (modeled as described above) and traits from other published studies, as described below. Significance values for correlations were determined and the false discovery rate controlled via the Benjamini and Hochberg method (Benjamini and Hochberg, 1995).

All traits analyzed for hierarchical clustering are divided into five major groups, as determined by the studies from which they are reported and by the phenotype that they measure, following the naming system described by Chitwood et al. (2013). “DEV” refer to leaf morphological and developmental traits as reported by Chitwood et al. (2013). “MET” traits report metabolite levels in the fruit pericarp, as measured by Schauer et al. (2006; 2008). Traits labeled “MOR” as recorded in Schauer et al. [2006; 2008] and Phenom-Networks include traits relevant to yield and morphological measures of fruits and flowers. The “ENZ” traits measure enzymatic activities in the fruit pericarp, as reported by Steinhauser et al. (2011), and “SEED” traits measure metabolite levels in seeds, as derived from Toubiana et al. (2012). Traits described in the present study are termed “WAX” because they relate to leaf waxes.

## 3 Results

### 3.1 Leaf wax traits from *S. pennellii* and *S. lycopersicum* cv M82 plants

Odd-carbon-numbered *n*-alkanes in *S. pennellii* and cv M82 ranged from C_27_ to C_35_, with C_31_ being the most abundant, followed by C_33_ (Table 1; Figure 2). Branched alkanes with methyl groups at the *iso* and *anteiso* positions are present in measurable quantities among both *S. pennellii* and cv M82.

**Figure 2:**
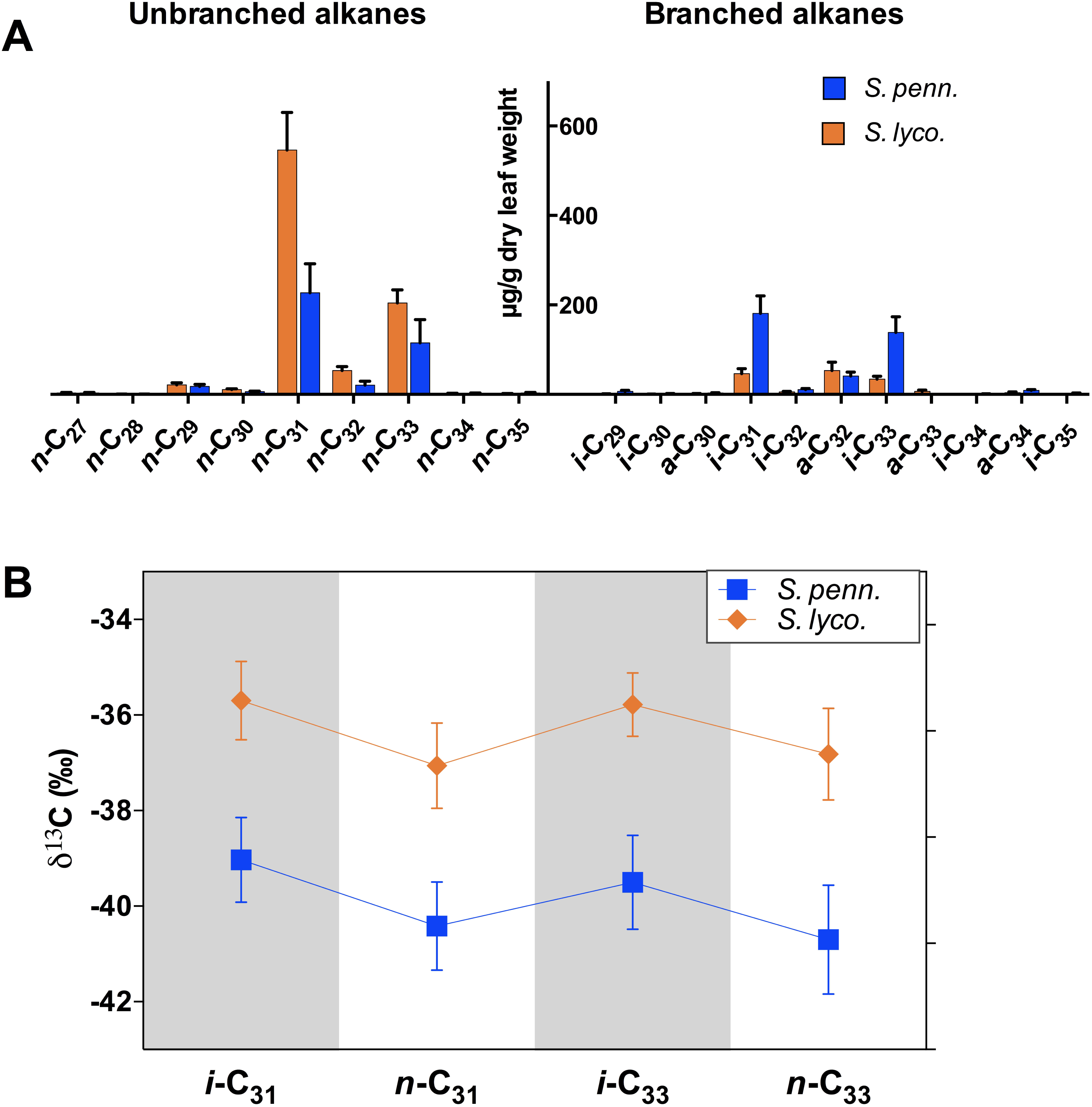
Leaf waxes from *S. lycopersicum* cv M82 and *S. pennellii.* **(A)** Concentration of leaf wax molecules, with unbranched (normal, left) and branched (right) alkanes. Average concentration values (pg/g leaf dry mass) of ten biological replicates are shown with error bars showing one standard deviation of the mean. *S. pennellii* (blue) contains much higher amounts of *iso*-alkanes than *S. lycopersicum* (orange). **(B)** Carbon isotope values (δ^13^C) measured from ten biological replicates analyzed in triplicate; analytical uncertainty ± 0.2 ‰, error bars represent standard deviation of biological replicates. The pattern of *iso-* over *n*-alkane enrichment is consistent between both species. Additionally, δ^13^C values from *S. pennellii* (blue) are consistently depleted relative to those from *S. lycopersicum* (orange).

**Table 1.** Leaf wax traits for *S. lycopersicum*, *S. pennellii*, and ILs. Average leaf wax trait values and standard deviations shown for the two parent lines grown simultaneously and for cv M82 grown with the ILs, and average trait values and ranges shown for IL plants. Broad sense heritability values (*H*^2^) shown as estimated for ILs. First column of data for *S. lycopersicum* and *S. pennellii* are from the first growth experiment; extra column of data for *S. lycopersicum* cv M82 and ILs are from the second growth experiment. * Indicates δ^13^C values reported.

| Traits | cv M82 | | *S. pennellii* | | cv M82<br>(with ILs) | | ILs | | $H^2$ |
| --- | --- | --- | --- | --- | --- | --- | --- | --- | --- |
|  | Mean | Stdev | Mean | Stdev | Mean | Stdev | Mean | Range |  |
| **Methylation index** | 0.15 | 0.02 | 0.50 | 0.04 | 0.13 | 0.01 | 0.14 | 0.07 to 0.28 | 0.35 |
| **Percent anteiso-alkanes** | 6.1 | 1.5 | 6.8 | 1.1 | 6.3 | 0.5 | 7.5 | 2.8 to 15.7 | 0.39 |
| **Percent**<br>***iso*-alkanes** | 8.8 | 0.7 | 43.6 | 2.6 | 6.5 | 0.7 | 7.0 | 3.3 to 13.1 | 0.37 |
| **CPI**<br>**[*n*-alkane]** | 10.4 | 0.4 | 11.2 | 1.3 | 11.8 | 0.8 | 12.5 | 9.6 to 17.4 | 0.31 |
| **CPI [*anteiso*-**<br>**alkane]** | 0.0 | 0.1 | 0.0 | 0.0 | 0.2 | 0.1 | 0.2 | 0.0 to 0.4 | 0.27 |
| **CPI**<br>**[*iso*-alkane]** | 10.2 | 0.6 | 19.4 | 2.5 | 9.6 | 1.7 | 12.3 | 5.8 to 21.2 | 0.22 |
| **ACL**<br>**[*n*-alkane]** | 31.5 | 0.0 | 31.5 | 0.1 | 31.3 | 0.1 | 31.3 | 31.0 to 31.8 | 0.30 |
| **ACL**<br>**[*anteiso*-**<br>**alkane]** | 32.1 | 0.0 | 32.2 | 0.0 | 32.1 | 0.1 | 32.1 | 31.9 to 32.3 | 0.28 |
| **ACL**<br>**[*iso*-alkane]** | 31.8 | 0.1 | 31.8 | 0.1 | 31.9 | 0.1 | 31.8 | 31.1 to 32.3 | 0.26 |
| **$\delta^{13}\text{C}(*)$ or $^{13}\epsilon$**<br>***i*-C<sub>31</sub> (‰)** | -35.7* | 0.8 | -39.0* | 0.9 | 16.7 | 2.0 | 18.2 | 13.1 to 22.0 | 0.18 |
| **$\delta^{13}\text{C}(*)$ or $^{13}\epsilon$**<br>***n*-C<sub>31</sub> (‰)** | -37.0* | 0.9 | -40.4* | 0.9 | 18.5 | 2.4 | 20.0 | 14.5 to 24.5 | 0.18 |
| **$\delta^{13}\text{C}(*)$ or $^{13}\epsilon$**<br>***i*-C<sub>33</sub> (‰)** | -35.8* | 0.7 | -39.5* | 1.0 | 16.4 | 2.0 | 18.0 | 13.4 to 21.8 | 0.13 |
| **$\delta^{13}\text{C}(*)$ or $^{13}\epsilon$**<br>***n*-C<sub>33</sub> (‰)** | -36.8* | 1.0 | -40.7* | 1.1 | 17.9 | 2.2 | 19.5 | 13.9 to 24.3 | 0.18 |
| **$\delta_n - \delta_{iso}$ (C<sub>31</sub>)**<br>**(‰)** | -1.4 | 0.5 | -1.4 | 0.3 | -1.6 | 0.3 | -1.7 | -0.3 to -4.0 | 0.39 |
| **$\delta_n - \delta_{iso}$ (C<sub>33</sub>)**<br>**(‰)** | -1.2 | 0.4 | -1.2 | 0.5 | -1.4 | 0.3 | -1.4 | -0.2 to -3.3 | 0.32 |

#### 3.1.1 Alkane methylation

The *S. pennellii* leaves produced more branched alkanes with methyl groups at the *iso* (second) position (Figure 2A). The average methylation index values are 0.15 and 0.50 for *S. lycopersicum* cv M82 and *S. pennellii,* respectively (Table 1), indicating that *S. pennellii* has a greater proportion of branched:normal alkanes than cv M82. This difference is driven by the higher percentage of *iso*-alkanes in *S. pennellii* (43.6% versus 8.8% in cv M82). The percent of *anteiso*-alkanes is indistinguishable between the two species.

#### 3.1.2 Structural traits

The *n*-alkane distributions for cv M82 and *S. pennellii* have high CPI (10.4 and 11.2, respectively; Table 1) and identical ACL (31.5) values. The *anteiso-* (3-methyl) alkanes CPI values of 0.0 for both cv M82 and *S. pennellii,* and a predominant even-numbered alkane distribution (ACL = 32.1 − 32.2). The *iso* (2-methyl) alkanes have high CPI values (10.2 and 19.4 for cv M82 and *S. pennellii,* respectively) and identical ACL values (31.8).

#### 3.1.3 Carbon isotopes

The average δ^13^C values of the most abundant leaf wax alkanes are reported in Figure 2B and Table 1. Among both cv M82 and *S. pennellii,* the *n*-alkanes are depleted relative to the *iso*-alkanes of length C_31_ and C_33_ by −1.4%o and −1.2%o, respectively (Table 1). *S. pennellii* alkanes are consistently depleted in ^13^C relative to those of cv M82. Mass balance calculations for these four major alkanes, which comprise 84% of all alkane mass measured, indicate that the average δ^13^C values for carbon incorporated in the leaf waxes for cv M82 is −36.9% and −39.9% for *S. pennellii.*

### 3.2 Leaf wax traits from IL plants

#### 3.2.1 Alkane methylation

Methylation indices for the ILs range from 0.07 to 0.28 (Table 1 and Supplemental Figure 4), with an average value of 0.14. The percentages of *anteiso-* and *iso*-alkanes range from 2.8% to 15.7% and from 3.3% to 13.1%, respectively. No IL approaches the percent of *iso*-alkanes measured from *S. pennellii* (43.6%). However, many ILs have a higher percentage of *anteiso*-alkanes than both *S. pennellii* and cv M82.

#### 3.2.2 Structural traits

The CPI values for the *n*-alkanes range from 9.6 to 17.4, from 0.0 to 0.4 for the *anteiso*-alkanes, and from 5.8 to 21.2 for *iso*-alkanes (Supplemental Figure 5, Table 1). ACL values vary from 31.0 to 31.8 for the *n*-alkanes, but alternate between odd and even predominance among the *anteiso-* and isoalkanes, ranging from 31.9 to 32.3 and from 31.1 and 32.3, respectively (Supplemental Figure 6, Table 1).

#### 3.2.3 Carbon isotopes

The ambient CO_2_ δ^13^C values ranged from −19.3‰ to −14.1‰ during the IL growth period, with the average defined as −16.4‰. Apparent fractionation, ^13^ε, is calculated according to Equation 2. Alkanes from the ILs vary in apparent fractionation compared to the alkanes from cv M82 (Table 1), ranging in ^13^ε values from 13.1‰ to 22.0‰ (*i*-C_31_), from 14.5% to 24.5% (*n*-C_31_), 13.4‰ to 21.8‰ (*i*-C_33_), and from 13.9‰ to 24.3‰ (*n*-C_33_). The ILs maintain the same pattern of carbon isotopic depletion of *n*-over *iso*-alkanes as measured from the parent alkanes. The magnitude of the depletion (δ_*n*_ - (δ_*iso*_) varies from −0.3 to −4.0 for C_31_ alkanes and from −0.2 to −3.3 for C_33_ alkanes.

### 3.3 Heritability and detected QTL

We modeled the suite of leaf wax traits measured from the ILs to estimate how much genetic factors play a role in differences in these traits between the IL plants and cv M82. Broad-sense heritability values (*H*^2^; Table 1 and Figure 2) for the leaf wax traits range from low (*H*^2^ = 0.13) to intermediate (0.39). The percentage of *anteiso-* (*H*^2^ = 0.39) and *iso*-alkanes (0.37) are of intermediate heritability, as are the magnitudes of carbon isotopic depletion of n-over *iso*-alkanes for C_31_ (0.39) and C_33_ (0.32). The methylation index (*H*^2^ = 0.35) is also of intermediate heritability. Traditional structural traits are also of intermediate heritability: CPI for *n*-alkanes (*H*^2^ = 0.31), *anteiso*-alkanes (0.27), and *iso*-alkanes (0.22), as well as ACL for *n*-alkanes (0.31), *anteiso*-alkanes (0.28), and *iso*-alkanes (0.26). Only the ^13^ε values of individual alkane molecules are of low heritability (*H*^2^ = 0.19 for *i*-C_31_, 0.18 for *n*-C_31_, 0.18 for *n*-C_33_, and 0.13 for *i*-C_33_).

We identified 156 QTLs at a significance level *p* < 0.05 for 14 leaf wax summary traits in this study (Figure 4, Supplemental Table 1); no QTL were detected for the carbon-isotopic enrichment of iso-over *n*-alkanes for C_33_. 124 QTL are significant in the direction toward *S. pennellii.* The QTL are determined relative to cv M82 grown simultaneously with the ILs.

#### 3.3.1 QTL regulating alkane methylation

QTL analysis may help to explain the genetic basis of variation in alkane methylation between *S. pennellii* and cv M82. *S. pennellii* has a greater methylation index (0.50) than any IL (varies from 0.07 to 0.28; Table 1). Ten QTLs of methylation index are significant in the direction of *S. pennellii,* i.e. greater than the methylation index of cv M82 (Figure 4, Supplemental Table 1, Supplemental Figure 4), which may be attributed to the variation induced by genetic loci of the ILs. There are eight QTL significant in the *S. pennellii* direction for percentages of *iso*-alkanes (Figure 4, Supplemental Table 1), whereas one QTL is transgressive beyond cv M82 (IL 8-1). Despite the small difference in the percent of *anteiso*-alkanes between *S. pennellii* and cv M82 (Table 1), there are eleven QTLs significant toward *S. pennellii* and one QTL that is transgressive beyond cv M82 (IL 7-5-5) (Figure 4, Supplemental Table 1).

Some ILs are identified as QTL for multiple alkane methylation traits. IL 3-2 displays the most significant increase in branched alkane production across three biological replicates, with an average methylation index of 0.26, 10.2% *iso*-alkanes, and 11.3% *anteiso*-alkanes. IL 1-1-2 has an average methylation index of 0.18, 8.9% *iso*-alkanes, and 9.3% *anteiso*-alkanes. From IL 1-1-3, we measured a methylation index of 0.20 and 10.5% *iso*-alkanes. IL 9-3-2 has a methylation index of 0.18 and 10.9% *anteiso*-alkanes. IL 7-4 has a methylation index of 0.18 and 9.7% *anteiso*-alkanes. ILs 4-3, 6-4, and 10-3 have methylation indices of 0.17, and 11.8%, 9.0%, and 9.5% *anteiso*-alkanes, respectively.

#### 3.3.2 QTL regulating CPI

Many QTL have been identified for CPI values: seven for *n*-alkanes, seven for *anteiso*-alkanes, and nineteen for *iso*-alkanes (Figure 4, Supplemental Table 1). For *n*-alkanes, the CPI values of identified QTL (between 13.5 to 16.2) are significant toward *S. pennellii.* Among *anteiso*-alkanes, the CPI values of identified QTL are all transgressive beyond cv M82. For the *iso*-alkanes, the identified QTL have CPI values ranging from 13.3 to 18.8 and are significant in the direction of *S. pennellii.*

#### 3.3.3 QTL regulating ACL

Thirty-nine QTLs have been identified for the three types of ACL values: seven for *n*-alkanes, seventeen for *anteiso*-alkanes, and fifteen for *iso*-alkanes (Figure 4, Supplemental Table 1). Among ACL values for *n*-alkane, three QTLs are significant in the direction of *S. pennellii* and four QTLs are transgressive beyond cv M82. For the *anteiso-alkane* ACL values, two QTLs are significant toward *S. pennellii,* whereas the remaining 15 are transgressive beyond cv M82. All fifteen of the iso-alkane ACL QTLs are significant in the direction of *S. pennellii.*

#### 3.3.4 QTL regulating carbon isotopic fractionation

We identified 54 QTLs for the five carbon isotope traits measured in this study: sixteen for ^13^ε *i*-C_31_ values, eleven for ^13^ε *i*-C_33_, ten for ^13^ε *n*-C_31_, eleven for ^13^ε *n*-C_33_, six for δ_*n*_ − δ_*iso*_ (C_31_), and zero for δ*n* − δ_*iso*_ (C_33_), (Figure 4, Supplemental Table 1). For all of the measured ^13^ε values, all QTLs are significant in the direction of *S. pennellii* and many QTLs overlap across all ^13^ε values: IL 1-1, 2-1, 3-5, 8-3-1, 9-3, 9-3-1, and 12-4-1. For the six QTLs of δ_*n*_ − δ_*iso*_ (C_31_), two are significant toward *S. pennellii*.

### 3.4 Hierarchical clustering

#### 3.4.1 Clustering of leaf wax traits

The clustering of leaf wax traits reveals similarities between the sets of trait measurements (Figure 5; Supplemental Dataset 7). Hierarchical clustering of wax traits groups the apparent fractionation values (^13^ ε) together. All of the ACL traits (*n*-, *iso,* and *anteiso*-alkanes) cluster together. CPI values for *iso*-alkanes cluster with the percent of *iso*-alkanes, which co-cluster with the CPI of *n*-alkanes. The carbon isotopic differences between *n*- and *iso*-alkanes (δ_*n*_ − δ_*iso*_) cluster with each other. The methylation index and percent of *anteiso*-alkanes are clustered together, and co-cluster with the CPI of *anteiso*-alkanes.

Numerous structural leaf wax traits are significantly correlated with each other based on multiple test-adjusted *p*-values and Spearman's ρ correlation coefficient. Among the CPI values, CPI of *anteiso*-alkanes correlates negatively with the CPI of *iso*-alkanes (*p* = 0.047, ρ = −0.28). The CPI values of *iso*-alkanes correlate positively with the percent of *iso*-alkanes (*p* = 0.002, ρ = 0.41). The CPI values of *n*-alkanes is negatively correlated with each of the methylation traits (methylation index, percent of *iso-* and *anteiso*-alkanes). The ACL of *n*-alkanes is positively correlated with *iso-* (*p* = 0.00, ρ = 0.82) and *anteiso*-alkanes (*p* = 0.00, ρ = 0.50). The ACL of *iso-* and *anteiso*-alkanes are negatively correlated with the percent of *anteiso*-alkanes. All ACL values are negatively correlated with the percent of *anteiso*-alkanes: with ACL values of *n*-(*p* = 0.025, ρ = −0.31), *iso-* (*p* = 0.003, ρ = −0.39), and *anteiso*-alkanes (*p* = 0.027, ρ = −0.30).

The methylation index is positively correlated with the percent of *iso-* and *anteiso*-alkanes and with the values of δ_*n*_ − δ_*iso*_ for both C_31_ and C_33_, but negatively correlated with ACL of *iso*-alkanes. The percent of *anteiso*-alkanes correlates positively with δ_*n*_ − δ_*iso*_ values of C_31_ and negatively with δ_*n*_ − δ_*iso*_ values of C_33_. The Δ values of all alkanes correlate positively with the CPI of *iso*-alkanes, and the ^13^ε values of all alkanes except *n*-C_33_ correlate positively with the percent of *iso*-alkanes. Most of the carbon isotopic traits significantly correlate with each other, except for δ_*n*_ − δ_*iso*_ with ^13^ε *i*− C_31_ and ^13^ε *i*-C_33_. The Δ values are positively correlated with each other; the ^13^ε values of *n*-C_31_ and *n*-C_33_ are positively correlated with the δ_*n*_ − δ_*iso*_ values of C_31_, but negatively correlated with the δ_*n*_ − δ_*iso*_ values of C_33_ (see Supplemental Dataset 7 for *p* and ρ values).

#### 3.4.2 Clustering of leaf wax traits with traits from previous studies

The traits measured in previous studies that cluster with WAX traits (Figure 6; Supplemental Figure 3) may contain information about the relevance of leaf wax traits to plant metabolism, yield, and developmental leaf traits. The CPI values for *n*-alkanes cluster with the enzymatic activity of succinyl-coenzyme A ligase in the fruit pericarp, and co-clusters with levels of uracil in the fruit pericarp and with activity levels of starch in the fruit pericarp (Schauer et al., 2006, 2008; Steinhauser et al., 2011). The percentage and CPI values of *iso*-alkanes co-cluster with enzymatic activity of invertase and glucokinase in the fruit pericarp (Steinhauser et al., 2011).

Multiple leaf wax traits from this study significantly correlate with traits measured in previous *S. pennellii* IL studies (Figure 6; Supplemental Dataset 8). ACL values for *n*- and *iso*-alkanes positively correlate with enzymatic activities of glyceraldehyde 3-phosphate dehydrogenase (GADPH; for *n*-alkane ACL, *p* = 0.013, ρ = 0.39; for *iso*-alkane ACL, *p* = 0.039, ρ = 0.34; Steinhauser et al., 2011) within the fruit pericarp, which serves as a catalyst during glycolysis. *iso*-Alkane ACL values positively correlate with metabolite levels of glutamate within seeds (*p* = 0.033, ρ = 0.36; Toubiana et al., 2012), an amino acid used to synthesize proteins. ACL values for *anteiso*-alkanes positively correlate with aconitase (*p* = 0.040, ρ = 0.34) and Suc phosphate synthase (*p* = 0.044, ρ = 0.34) enzymatic activity within the fruit pericarp (Steinhauser et al., 2011). CPI values for *iso*-alkanes positively correlate with the size of epidermal pavement cells (*p* = 0.033, ρ = 0.36; Chitwood et al., 2013), which form a protective layer for more specialized cells on leaves, and negatively correlate with fructose levels in the fruit pericarp (*p* = 0.033, ρ = −0.35; Steinhauser et al., 2011) and with the levels of metabolite trehalose within seeds (*p* = 0.022, ρ = −0.39; Toubiana et al., 2012).

Multiple flower morphological traits correlate positively with δ_*n*_− δ_*iso*_ for C_31_: anther length (*p* = 0.041, ρ = 0.34), measures of the ratio of style length:width (*p* = 0.009, ρ = 0.40), and style length (*p* = 0.001, ρ = 0.49). Conversely, these same traits negatively correlate with δ_*n*_− δ_*iso*_ for C_33_: anther length (*p* = 0.006, ρ = −0.42), measures of the ratio of style length:width (*p* = 0.022, ρ = −0.37), and style length (*p* = 0.003, ρ = −0.44) (Schauer et al., 2006, 2008). δ_*n*_− δ_*iso*_ for C_31_ correlates positively with levels of benzoate in the seeds (p = 0.032, ρ = 0.36; Toubiana et al., 2012) and negatively with enzymatic activities of phosphofructokinase a (p = 0.039, ρ = −0.34; Steinhauser et al., 2011), which is involved in sugar metabolism. δ_*n*_− δ_*iso*_ for C_33_ correlates negatively with measures of fruit width (p = 0.038, ρ = −0.35) and weight (p = 0.012, ρ = −0.39), and with the weight of the seeds (p = 0.016, ρ = −0.368; Schauer et al., 2006, 2008). The values of δ_*n*_− δ_*iso*_ for C_33_ correlate positively with the metabolic activity of fumarate in the fruit pericarp (*p* = 0.015, ρ = 0.39; Schauer et al., 2006, 2008) and with the levels of adenine within seeds (*p* = 0.046, ρ = 0.35; Toubiana et al., 2012). The number of flowers per inflorescence correlates negatively with ^13^ε values for *i*-C_31_ (*p* = 0.011, ρ = −0.42; Schauer et al., 2006, 2008).

## 4 Discussion

Results from QTL analysis may help to explain the genetic basis of variation in leaf wax traits between *S. pennellii* and cv M82. As evidenced by the multiple QTL identified for nearly all leaf wax traits in this study (except for δ_*n*_− δ_*iso*_ of C33; Figure 4, Supplemental Table 1), large portions of the genome contribute to natural variation in many leaf wax traits, suggesting that these traits are polygenic.

### 4.1 Alkane methylation

No IL has a methylation index or percent of *iso*-alkanes comparable to those of *S. pennellii* (Table 1). All methylation traits are of intermediate heritability (Figure 3), indicating that these traits are moderately influenced by genetic controls. ILs 1-1-2, 1-1-3, 1-4, and 3-2 are among the greatest contributing loci to alkane methylation, with significant QTL for both methylation indices and percentages of *iso*-alkanes (Figure 4; Supplemental Table 1). The variation among the QTL significant for methylation traits may be attributed to the variation induced by the genetic loci of the ILs.

**Figure 3:**
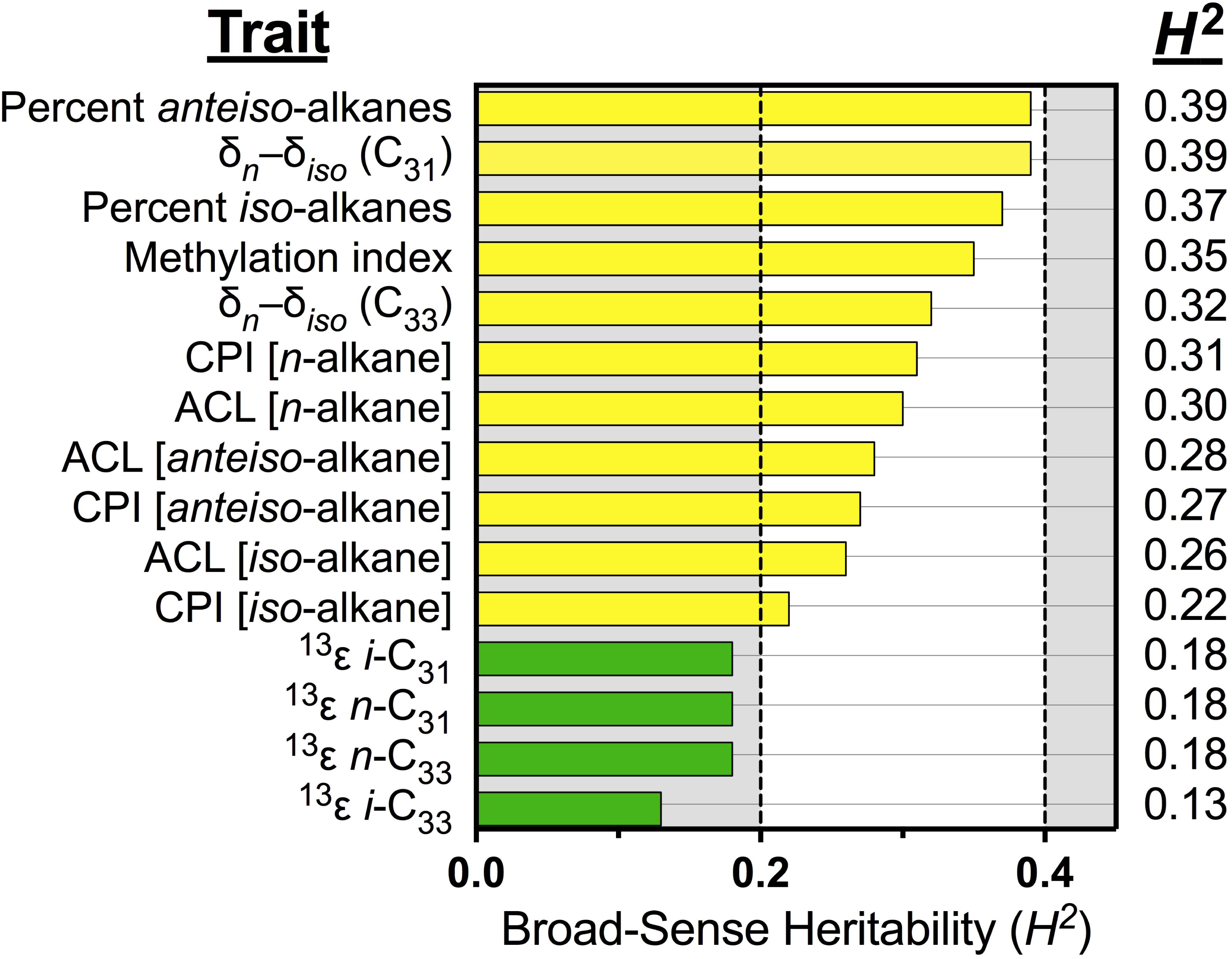
Broad-sense heritability for leaf wax traits. Colors denote traits with intermediate (yellow, 0.2 ≤ *H^2^* < 0.4), and low (green, *H^2^* < 0.2) heritability values. All traits were measured from plants grown under 2014 St. Louis, MO greenhouse conditions. The majority of leaf wax traits have intermediate heritability values. CPI measured from *anteiso*-alkanes and individual carbon isotope values have low heritability values.

**Figure 4:**
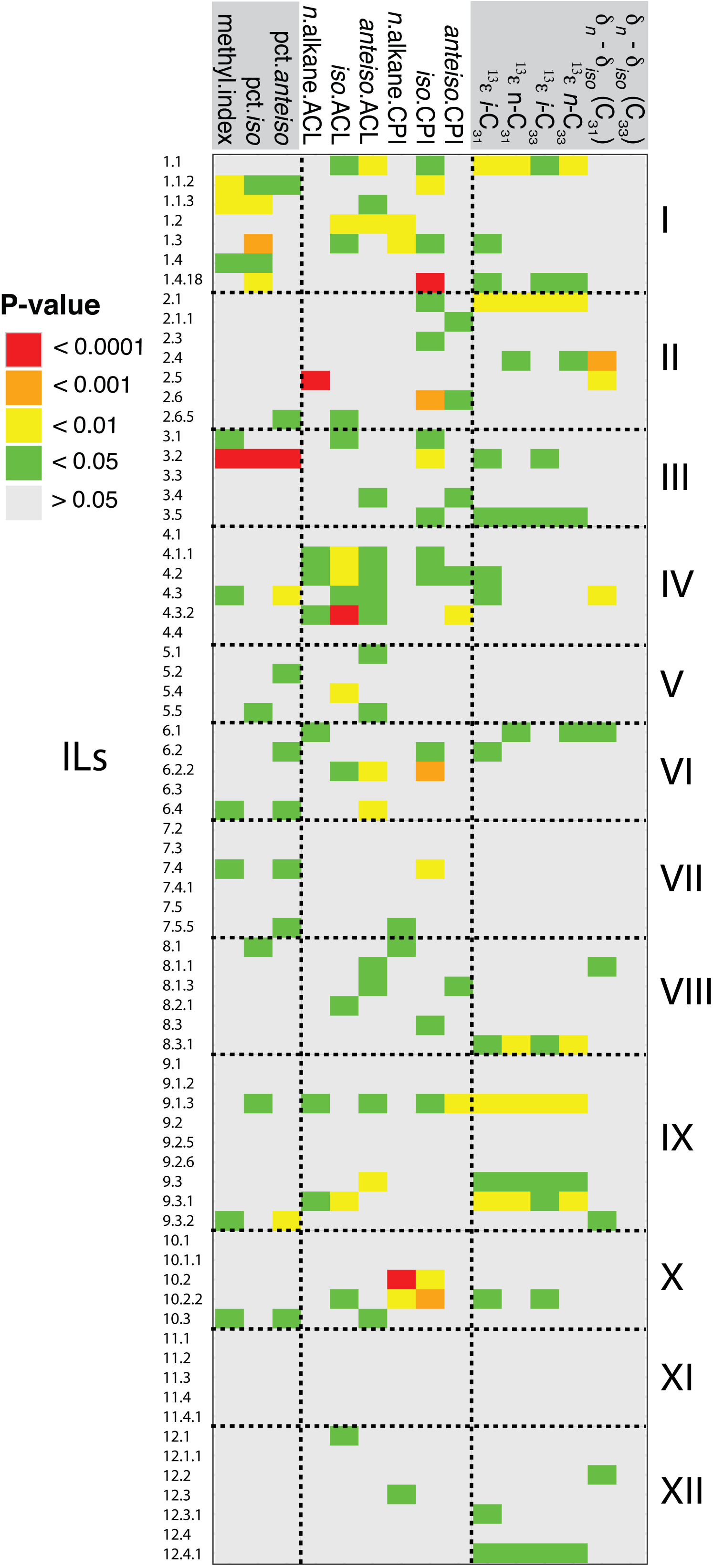
Detected leaf wax QTLs. Shown are QTL with *p*-value < 0.05, as calculated from mixed-effect linear models for deviation of ILs from cv M82. In total, 139 QTLs were detected. Traits are grouped by type: from left to right, structural, carbon isotope, and methylation traits. White spaces represent traits for which no IL replicates had quantifiable data.

We observed many significant correlations between methylation traits and other leaf wax traits (Figure 5; Supplemental Dataset 7). Among the ILs, the methylation traits decrease with increasing ACL for any type of alkane and with CPI of *n*- and *anteiso*-alkanes. However, methylation traits increase with increasing CPI values of *iso*-alkanes.

**Figure 5:**
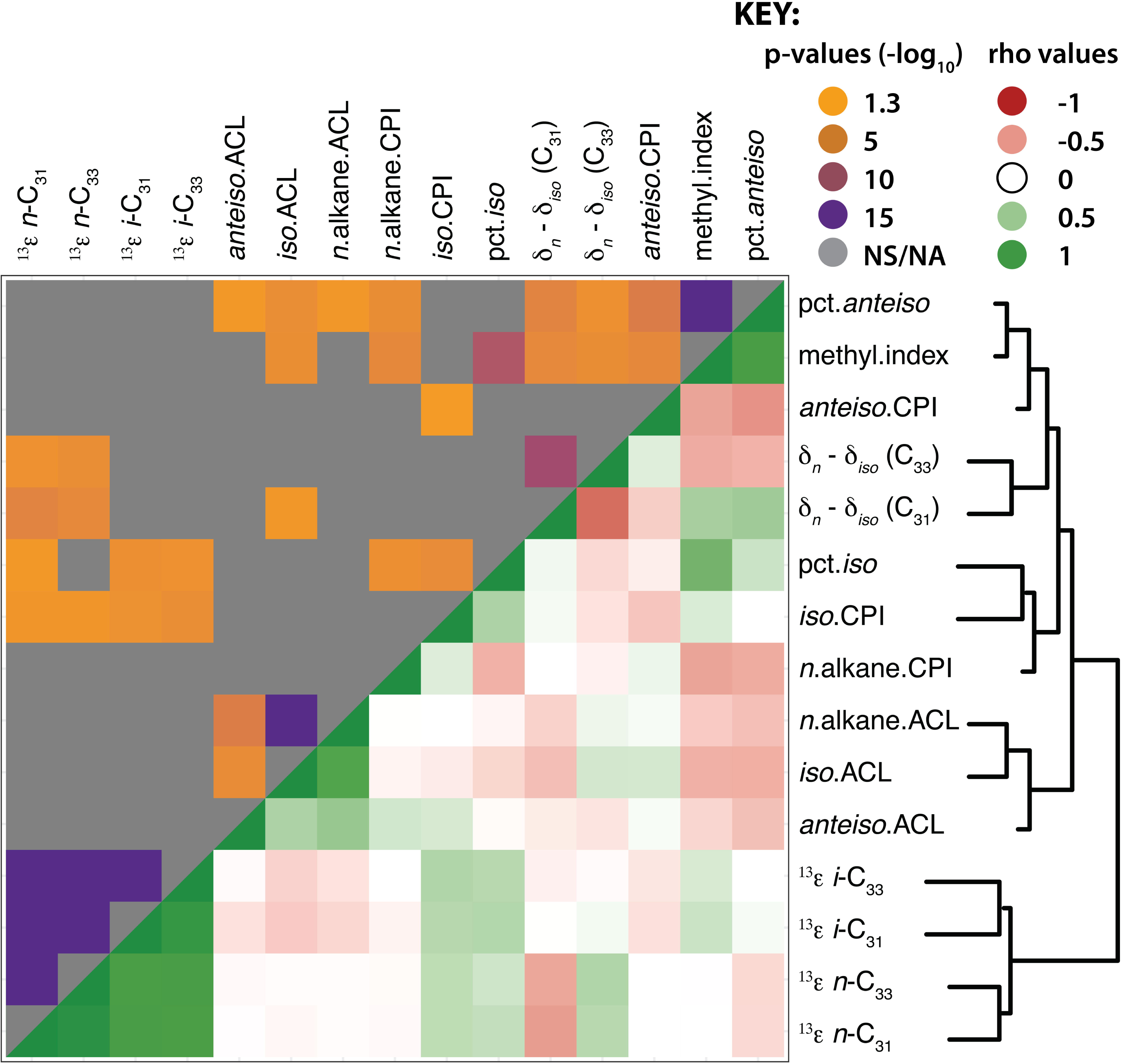
Hierarchical clustering and correlation of leaf wax traits. Hierarchical clustering and heat map of leaf wax traits measured in this study with each other. The upper quadrant shows correlation *p*-values; gray indicates non-significant *p*-values (*p* > 0.05), whereas the spectrum of orange to purple colors designate *p*-values ranging from less to more significant, respectively. The lower quadrant indicates Spearman's ρ values in red (negative), white (neutral), and green (positive).

### 4.2 Structural traits

The prevalence of *n*-C_31_ among the leaves of cv M82 and *S. pennellii* in this study is identical to that previously reported in the fruit cuticles of the same plants (Yeats et al., 2012). Reddy et al. (2000) measured that *iso*-alkanes are predominantly odd-numbered and that *anteiso*-alkanes are predominantly even-numbered among *Micromeria* plants, which is the same pattern noted for plants in this study. Among the *Micromeria* plants, the CPI values range from 5.6 to 7.2 for *n*-alkanes, from 0.20 to 0.34 for *anteiso*-alkanes, and from 3.9 to 5.3 for *iso*-alkanes (Reddy et al., 2000). Compared to *S. pennellii* and cv M82 in this study, the CPI values for *anteiso*-alkanes from *Micromeria* are of similar magnitude, whereas the CPI values for n- (from 9.6 to 17.4; Table 1) and *iso*-alkanes (5.8 to 21.2) have a greater range among the ILs.

Numerous studies have revealed correlations between *n*-alkane chain lengths and climatic variables such as temperature and humidity, as well as to the plant sources of the *n*-alkanes (e.g., Bush and McInerney, 2013 and references therein). Given this correlation, we might expect ACL values to be different in plants that are adapted to different hydrological regimes (e.g., *S. pennellii* and cv M82) and low heritability of ACL traits. Instead, we observe nearly identical ACL values between *S. pennellii* and cv M82 (Table 1) and ACL values that are of intermediate heritability (Figure 3), suggesting a high degree of genetic control over their alkane chain-length distributions.

A benefit to studying the *S. pennellii* IL library is the ability to correlate phenotypic data sets across multiple growth studies in order to probe how leaf waxes relate to other phenotypes measured from the fruit and leaves of the same genetic variants. Our hierarchical clustering analysis reveals that ACL values for *anteiso*-alkanes positively correlate with aconitase enzymatic activity within the fruit pericarp (Figure 6; Supplemental Dataset 8; Steinhauser et al., 2011), which might be related to *anteiso-alkane* synthesis. Grice et al., (2008) propose that the methylbutyryl-CoA moiety derived from isoleucine is the precursor molecule for *anteiso*-alkanes. Oxaloacetate is the precursor for isoleucine synthesis. Aconitase catalyzes the isomerization of citrate to isocitrate, which can be cyclically decarboxylated into oxaloacetate for export to the chloroplast and used for isoleucine synthesis. Grice et al. (2008) suggest that isoleucine sourced from isocitrate might be isotopically heavy because it is sourced from the cytosol; in the present study, *anteiso*-alkanes are not abundant enough to make carbon isotope measurements. In our study, leaf wax traits do not correlate with levels of isoleucine or isocitrate measured from fruit pericarp or with isoleucine abundances measured in seeds.

**Figure 6:**
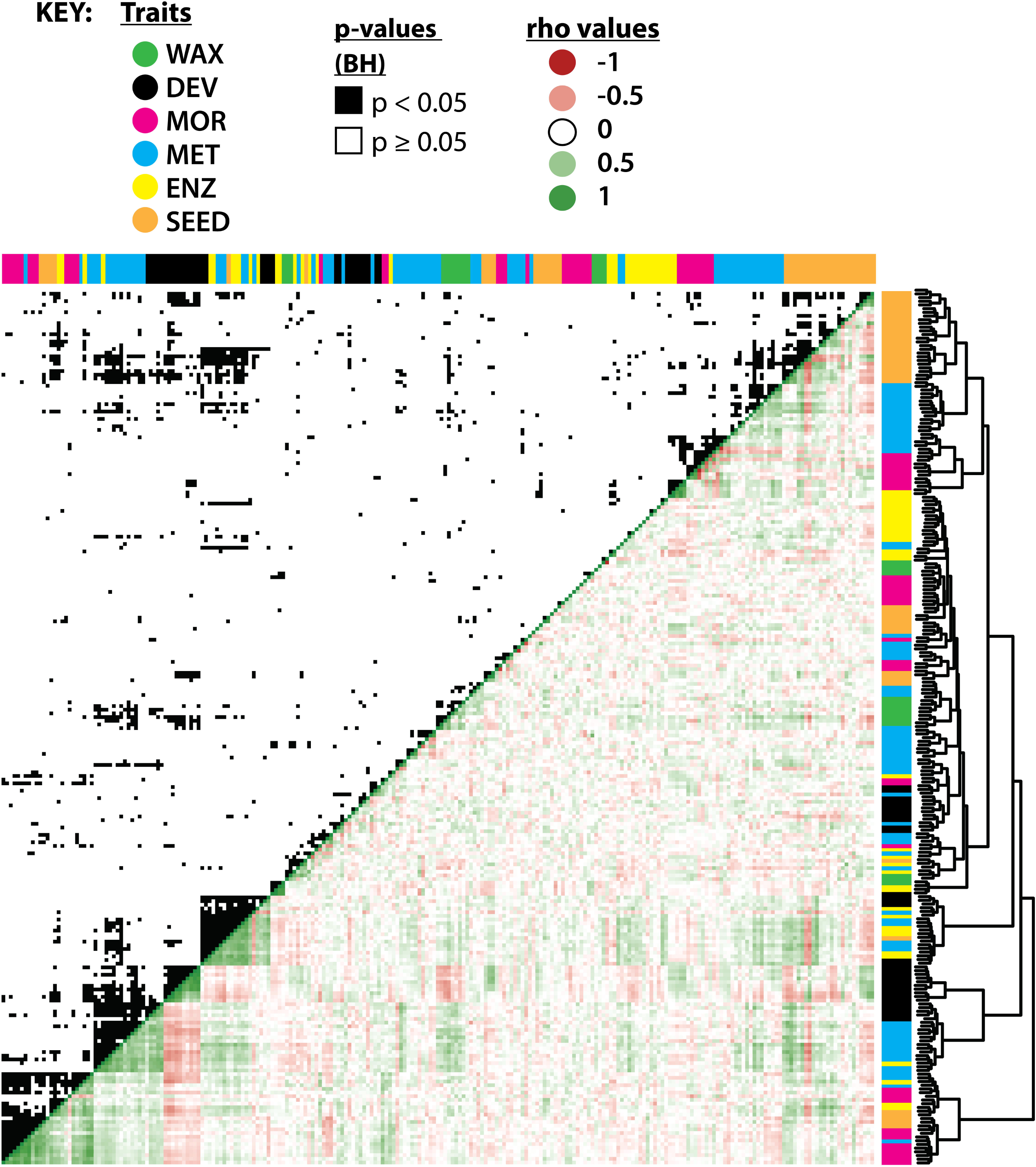
Hierarchical clustering and correlation of leaf wax traits with previously measured IL traits. Hierarchical clustering of leaf wax traits from this study with traits measured in previous studies (see close-up of the hierarchical clustering in Supplemental Figure 3). WAX (green) are leaf wax traits from this study; DEV (black), leaf developmental traits from Chitwood *et al.* (2013); MOR (pink), entire-plant, yield, and reproductive morphological traits from Schauer *et al.* (2006, 2008); MET (blue), metabolic traits from the two previous studies; ENZ (yellow), enzymatic activities from Steinhauser *et al.* (2011); SEED (orange), seed metabolites as measured by Toubiana *et al.* (2012). Hierarchical clustering is based on absolute correlation values. The upper quadrant shows significant correlations (*p* < 0.05) between traits after multiple test adjustment, shown in black. The lower quadrant indicates Spearman's ρ values in red (negative), white (neutral), and green (positive)

### 4.3 Carbon isotopes

The ^13^ε values for the four primary alkanes in this study have low broad-sense heritability values (Figure 3). The low heritability reflects the significant isotopic variation among biological replicates. A previous study into the bulk carbon isotopic composition of *Arabidopsis thaliana* grown in controlled growth chambers measured high heritability for bulk leaf δ^13^C values (*H^2^* = 0.67; Easlon et al., 2014). The lower heritability found among δ^13^C of wax in this study may reflect the more variable environmental conditions of a greenhouse relative to a growth chamber, or a biological difference between *Arabadopsis* and *Solanum.* Broad-sense heritability is specific to the population and environment, thus the difference among results is not unexpected.

We observe that there is an intrinsic biological difference in δ^13^C values between *S. pennellii* and cv M82: *S. pennellii* alkanes are consistently depleted by roughly 3%o in ^13^C relative to cv M82 (Figure 2). Although the δ^13^CCO_2_ values within the greenhouse varied by at least 3‰ during the growth period of the parent lines (Supplemental Figure 2; Supplemental Dataset 3), the offset in δ^13^C values likely does not result from different timing of carbon fixation between the plants, given that we sampled contemporaneous leaf material from all plants.

To explore the correlation between water use efficiency (which is estimated by carbon isotope composition) and stomatal conductance (Easlon et al., 2014), we tested our hierarchical clustering analysis for correlations between leaf stomatal density measurements made by Chitwood et al. (2013) and our leaf wax carbon isotope traits; however, our analysis yielded no significant correlations.

Among all plants in this study, *iso*-alkanes are enriched in ^13^C over *n*-alkanes, expressed here as δ_*n*_−δ_*iso*_. These traits are of intermediate heritability (*H*^2^ = 0.38 for C_31_ alkanes and *H*^2^ = 0.32 for C_33_ alkanes; Figure 3), suggesting that the enrichment is strongly influenced by genetic controls. It is interesting that the isotopic enrichment is more heritable than individual carbon isotopic measurements. Reddy et al. (2000) noted no apparent differences in δ^13^C values between normal and branched alkanes in their study of four species of *Micromeria.* However, the enrichment pattern observed in this study is consistent with that reported by Grice et al. (2008), who recorded that *iso*-alkanes are enriched by 0-1.8%o over *n*-alkanes in *Nicotiana tabacum* (tobacco) plants. Grice et al. (2008) attributed this enrichment pattern to different biosynthetic precursors for *iso-* versus *n*-alkanes (i.e., valine for *iso-* and pyruvate for *n*-alkanes). Levels of valine and pyruvate have previously been measured from both the fruits (Schauer et al., 2006; 2008) and seeds (Toubiana et al., 2012) of the *S. pennellii* ILs; however, these traits do not significantly correlate with any WAX traits in this study.

## 5 Implications for interpreting sedimentary plant waxes

Although *S. lycopersicum* cv M82 and *S. pennellii* are not abundant producers of leaf waxes in terrestrial ecosystems, they nonetheless provide a useful tool for investigating plant genetics and physiology. We demonstrate in this study that the use of this model species complex allows us to determine the role of genetic versus environmental influences in the production of plant wax traits that are central to paleoclimatic reconstruction. This approach allows us to test the implicit assumptions in paleoclimate applications about the importance, or lack thereof, of genetic influence over leaf wax paleoclimate proxies.

Carbon isotope values (δ^13^C) of plant materials from sediments can be used to identify ecosystems dominated by C_3_ versus C_4_ plants. Among individual plants, δ^13^C is positively correlated with water use efficiency of plants (e.g., Easlon et al., 2014), which can plastically respond to changing local rainfall and humidity. An implicit assumption for using δ^13^C values to interpret changes in water use efficiency is that the δ^13^C alkane signal is dominated by environmental rather than genetic information. By examining this assumption, we have quantified that the δ^13^C values of leaf waxes measured from plants in this study are strongly influenced by environmental variance (*H*^2^ ranges from 0.13 to 0.19). Our study reveals that genetic variance plays a limited role in driving variation of leaf wax carbon isotopic values among *Solanum* plants, and is consistent with the interpretation that variation in the δ^13^C of wax compounds, as recorded in sediments, is largely driven by paleohydrological changes. These findings do not bear on changes in the input of plants with a strongly different carbon fixation pathway, such as C_4_ plants.

Given the correlations between *n*-alkane chain lengths and climatic variables such as temperature and humidity, we might expect ACL values to be strongly influenced by environmental cues. Rather, we measure ACL values that are of intermediate heritability (0.30), suggesting a strong degree of genetic influence over alkane chain-length distributions. Future studies that utilize this model species complex in different environments might further illuminate the connection between alkane distributions and climatic variables.

All alkane methylation traits in this study are largely influenced by genetic variation, which is in agreement with the fact that branched alkanes have been identified from only a few modern plant families (see Introduction). The presence of branched alkanes in the sedimentary record might lend itself to chemotaxonomic applications, but it is unlikely that any of the branched alkane-producing plants are significant global contributors to terrestrial soil organic matter. However, regional chemotaxonomic applications of branched alkanes have proved useful. For example, Pautler et al. (2014) identified that an Arctic chickweed contributed to sedimentary organic matter based on the presence of branched alkanes. Fukushima et al. (2005) used the presence of *anteiso* compounds to suggest a local proxy for lake acidification. Branched alkanes and the methylation index can be more useful for chemotaxonomic applications on a regional level.

The *n*-alkane hydrogen isotope proxy (δ^2^H) is assumed to record environmental information with minimal complications introduced from genetic variability. Thus, the environmental controls are assumed to be dominant over phenotypic variability. A future report will present results that examine this assumption under controlled conditions using this set of model organisms, and thus quantify the relative proportions of genetic and environmental influences over leaf wax δ^2^H values.

## 6

### Acknowledgements

From assisting with plant collections to helping with isotopic analyses, laboratory manager Melanie Suess (Washington University in St. Louis) was instrumental to this project. We thank undergraduate researchers Jenny Zhang and Claire Ma (Washington University in St. Louis) for their contributions. We thank postdoctoral researchers Margaret Frank and Viktoriya Koneva (Donald Danforth Plant Science Center) and research scientist Jen Houghton (Washington University in St. Louis) for their assistance during leaf collection. We also thank David Fike and Dwight McCay (Washington University in St. Louis) for loaning and assisting with setup of the Picarro instrumentation. We thank Kevin Reilly (Donald Danforth Plant Science Center) for caring for the plants in the greenhouse, and Dani Zamir (Hebrew University, Rehovot, Israel), the Tomato Genetics Resource Center, and the lab of Neelima Sinha (University of California, Davis) for gifts of germplasm.

## Funding

Acknowledgment is made to the Donors of the American Chemical Society Petroleum Research Fund for partial support of this research through PRF grant #53417-DNI2. Partial funding for this work was provided by I-CARES, Washington University in Saint Louis. We also acknowledge the support of Washington University in St. Louis and the Donald Danforth Plant Science Center.

## References

Abràmoff, M. D., Hospitals, I., Magalhães, P. J., and Abràmoff, M. (2004). Image Processing with ImageJ. Biophotonics Int. 11, 36–42.

Benjamini, Y., and Hochberg, Y. (1995). Controlling the False Discovery Rate: A Practical and Powerful Approach to Multiple Testing. J. R. Stat. Soc. Ser. B 57, 289–300.

Bolger, A., Scossa, F., Bolger, M. E., Lanz, C., Maumus, F., Tohge, T., et al. (2014). The genome of the stress-tolerant wild tomato species Solanum pennellii. Nat. Genet. 46, 1034–1038. doi:10.1038/ng.3046.

Bush, R. T., and McInerney, F. a. (2013). Leaf wax n-alkane distributions in and across modern plants: Implications for paleoecology and chemotaxonomy. Geochim. Cosmochim. Acta 117, 161–179. doi:10.1016/j.gca.2013.04.016.

Bush, R. T., and McInerney, F. A. (2015). Influence of temperature and C4 abundance on n-alkane chain length distributions across the central USA. Org. Geochem. 79, 65–73. doi: 10.1016/j.orggeochem.2014.12.003.

Castañeda, I. S., Mulitza, S., Schefuss, E., dos Santos, R. A. L., Damste, J. S. S., and Schouten, S. (2009a). Wet phases in the Sahara/Sahel region and human migration patterns in North Africa. Proc. Natl. Acad. Sci. U. S. A. 106, 20159–20163. doi:Doi 10.1073/Pnas.0905771106.

Castañeda, I. S., Werne, J. P., Johnson, T. C., and Filley, T. R. (2009b). Late Quaternary vegetation history of southeast Africa: The molecular isotopic record from Lake Malawi. Palaeogeogr. Palaeoclimatol. Palaeoecol. 275, 100–112. doi:10.1016/j.palaeo.2009.02.008.

Chitwood, D. H., Kumar, R., Headland, L. R., Ranjan, A., Covington, M. F., Ichihashi, Y., et al. (2013). A quantitative genetic basis for leaf morphology in a set of precisely defined tomato introgression lines. Plant Cell 25, 2465–81. doi:10.1105/tpc.113.112391.

Diefendorf, A. F., Freeman, K. H., Wing, S. L., and Graham, H. V. (2011). Production of n-alkyl lipids in living plants and implications for the geologic past. Geochim. Cosmochim. Acta 75, 7472–7485. doi:10.1016/j.gca.2011.09.028.

Diefendorf, A. F., Mueller, K. E., Wing, S. L., Koch, P. L., and Freeman, K. H. (2010). Global patterns in leaf 13C discrimination and implications for studies of past and future climate. Proc. Natl. Acad. Sci. 107, 5738–5743. doi:10.1073/pnas.0910513107.

Easlon, H. M., Nemali, K. S., Richards, J. H., Hanson, D. T., Juenger, T. E., and McKay, J. K. (2014). The physiological basis for genetic variation in water use efficiency and carbon isotope composition in Arabidopsis thaliana. Photosynth. Res. 119, 119–129. doi:10.1007/s11120-013-9891-5.

Eglinton, G., Gonzalez, A. G., Hamilton, R. J., and Raphael, R. A. (1962). Hydrocarbon Constituents of the Wax Coatings of Plant Leaves: A Taxonomic Survey. Phytochemistry 1, 89–102.

Eglinton, T. I., Eglinton, G., Dupont, L. M., Sholkovitz, E. R., Montluçon, D., and Reddy, C. M. (2002). Composition, age, and provenance of organic matter in NW African dust over the Atlantic Ocean. Geochemistry, Geophys. Geosystems 3, 27p. doi:10.1029/2001GC000269.

Eshed, Y., and Zamir, D. (1995). An Introgression Line Population of Lycopersicon pennellii in the Cultivated Tomato Enables the Identification and Fine Mapping of Yield-Associated QTL. Genetics 141, 1147–1162.

Farquhar, G. D., Ehleringer, J. R., and Hubick, K. T. (1989). Carbon Isotope Discrimination and Photosynthesis. Annu. Rev. Plant Physiol. Plant Mol. Biol. 40, 503–537. doi:1040-2519/89/0601-503.

Feakins, S. J., Eglinton, T. I., and deMenocal, P. B. (2007). A comparison of biomarker records of northeast African vegetation from lacustrine and marine sediments (ca. 3.40 Ma). Org. Geochem. 38, 1607–1624. doi:10.1016/j.orggeochem.2007.06.008.

Feakins, S. J., Peter, B., and Eglinton, T. I. (2005). Biomarker records of late Neogene changes in northeast African vegetation. Geology 33, 977–980. doi:10.1130/G21814.1.

Feakins, S. J., and Sessions, A. L. (2010). Controls on the D/H ratios of plant leaf waxes in an arid ecosystem. Geochim. Cosmochim. Acta 74, 2128–2141. doi:10.1016/j.gca.2010.01.016.

Fobes, J. F., Mudd, B. J., and Marsden, M. P. F. (1985). Epicuticular Lipid Accumulation on the Leaves of Lycopersicon pennellii (Corr.) D'Arcy and Lycopersicon esculentum Mill. Plant Physiol. 77, 567–570.

Fukushima, K., Yoda, A., Kayama, M., and Miki, S. (2005). Implications of long-chain anteiso compounds in acidic freshwater lake environments: Inawashiro-ko in Fukushima Prefecture, Japan. Org. Geochem. 36, 311–323. doi:10.1016/j.orggeochem.2004.07.014.

Futuyma, D. J. (1998). Evolutionary Biology. 3rd ed. Sunderland, Massachusetts: Sinauer Associates, Inc.

Gao, L., Edwards, E. J., Zeng, Y., and Huang, Y. (2014). Major evolutionary trends in hydrogen isotope fractionation of vascular plant leaf waxes. PLoS One 9. doi:10.1371/journal.pone.0112610.

Gao, L., Guimond, J., Thomas, E., and Huang, Y. (2015). Major trends in leaf wax abundance, δ2H and δ13C values along leaf venation in five species of C3 plants: Physiological and geochemical implications. Org. Geochem. 78, 144–152. doi:10.1016/j.orggeochem.2014.11.005.

Girard, A.-L., Mounet, F., Lemaire-Chamley, M., Gaillard, C., Elmorjani, K., Vivancos, J., et al. (2012). Tomato GDSL1 is required for cutin deposition in the fruit cuticle. Plant Cell 24, 3119–34. doi:10.1105/tpc.112.101055.

Grice, K., Lu, H., Zhou, Y., Stuart-Williams, H., and Farquhar, G. D. (2008). Biosynthetic and environmental effects on the stable carbon isotopic compositions of anteiso-(3-methyl) and iso-(2-methyl) alkanes in tobacco leaves. Phytochemistry 69, 2807–2814. doi:10.1016/j.phytochem.2008.08.024.

Heemann, V., Brümmer, U., Paulsen, C., and Seehofer, F. (1983). Composition of the leaf surface gum of some Nicotiana species and Nicotiana tabacum cultivars. Phytochemistry 22, 133–135. doi:10.1016/S0031-9422(00)80073-4.

Hou, J., D'Andrea, W. J., MacDonald, D., and Huang, Y. (2007a). Evidence for water use efficiency as an important factor in determining the δD values of tree leaf waxes. Org. Geochem. 38, 1251–1255. doi: 10.1016/j.orggeochem.2007.03.011.

Hou, J., D'Andrea, W. J., MacDonald, D., and Huang, Y. (2007b). Hydrogen isotopic variability in leaf waxes among terrestrial and aquatic plants around Blood Pond, Massachusetts (USA). Org. Geochem. 38, 977–984. doi:10.1016/j.orggeochem.2006.12.009.

Huang, X., Meyers, P. a., Wu, W., Jia, C., and Xie, S. (2011). Significance of long chain iso and anteiso monomethyl alkanes in the Lamiaceae (mint family). Org. Geochem. 42, 156–165. doi:10.1016/j.orggeochem.2010.11.008.

Jetter, R., Kunst, L., and Samuels, L. (2006). “Composition of plant cuticular waxes,” in Biology of the Plant Cuticle, 145–181.

Kosma, D. K., Bourdenx, B., Bernard, a., Parsons, E. P., Lu, S., Joubes, J., et al. (2009). The Impact of Water Deficiency on Leaf Cuticle Lipids of Arabidopsis. Plant Physiol. 151, 1918–1929. doi: 10.1104/pp. 109.141911.

Naafs, B. D. A., Hefter, J., Acton, G., Haug, G. H., Martínez-Garcia, A., Pancost, R., et al. (2012). Strengthening of North American dust sources during the late Pliocene (2.7Ma). Earth Planet. Sci. Lett. 317–318, 8–19. doi:10.1016/j.epsl.2011.11.026.

Pautler, B. G., Austin, J., Otto, A., Stewart, K., Lamoureux, S. F., Simpson, M. J., et al. (2014). Biomarker assessment of organic matter sources and degradation in Canadian High Arctic littoral sediments. Biogeochemistry 100, 75–87. doi:10.1007/sl0533-009-9405-x.

Polissar, P. J., and D'Andrea, W. J. (2014). Uncertainty in paleohydrologic reconstructions from molecular δD values. Geochim. Cosmochim. Acta 129, 146–156. doi:10.1016/j.gca.2013.12.021.

Reddy, C. M., Eglinton, T. I., Palić, R., Benitez-Nelson, B. C., Stojanović, G., Palić, I., et al. (2000). Even carbon number predominance of plant wax n-alkanes: A correction. Org. Geochem. 31, 331–336. doi:10.1016/S0146-6380(00)00025-5.

Rogge, W., and Hildemann, L. (1994). Sources of fine organic aerosol. 6. Cigarette smoke in the urban atmosphere. Environ. Sci. Technol. 28, 1375–1388. doi:10.1021/es00056a030.

Sachse, D., Billault, I., Bowen, G. J., Chikaraishi, Y., Dawson, T. E., Feakins, S. J., et al. (2012). Molecular Paleohydrology: Interpreting the Hydrogen-Isotopic Composition of Lipid Biomarkers from Photosynthesizing Organisms. Annu. Rev. Earth Planet. Sci. 40, 221–249. Available at: http://www.annualreviews.org/doi/abs/10.1146/annurev-earth-042711-105535.

Schauer, N., Semel, Y., Balbo, I., Steinfath, M., Repsilber, D., Selbig, J., et al. (2008). Mode of Inheritance of Primary Metabolic Traits in Tomato. Plant Cell 20, 509–523. doi:10.1105/tpc.107.056523.

Schauer, N., Semel, Y., Roessner, U., Gur, A., Balbo, I., Carrari, F., et al. (2006). Comprehensive metabolic profiling and phenotyping of interspecific introgression lines for tomato improvement. Nat. Biotechnol. 24, 447–454. doi:10.1038/nbt1192.

Schubert, B. A., and Jahren, A. H. (2012). The effect of atmospheric CO2 concentration on carbon isotope fractionation in C3 land plants. Geochim. Cosmochim. Acta 96, 29–43.

Silva, K. M. M. Da, Agra, M. D. F., Santos, D. Y. A. C. Dos, and Oliveira, A. F. M. De (2012). Leaf cuticular alkanes of Solanum subg. Leptostemonum Dunal (Bitter) of some northeast Brazilian species: Composition and taxonomic significance. Biochem. Syst. Ecol. 44, 48–52. doi:10.1016/j.bse.2012.04.010.

Smirnova, A., Leide, J., and Riederer, M. (2013). Deficiency in a very-long-chain fatty acid β-ketoacyl-coenzyme a synthase of tomato impairs microgametogenesis and causes floral organ fusion. Plant Physiol. 161, 196–209. doi:10.1104/pp.112.206656.

Smith, F. a., and Freeman, K. H. (2006). Influence of physiology and climate on δD of leaf wax n-alkanes from C3 and C4 grasses. Geochim. Cosmochim. Acta 70, 1172–1187. doi:10.1016/j.gca.2005.11.006.

Smith, R. M., Marshall, J. A., Davey, M. R., Lowe, K. C., and Power, J. B. (1996). Comparison of volatiles and waxes in leaves of genetically engineered tomatoes. Phytochemistry 43, 753–758.

Steinhauser, M.-C., Steinhauser, D., Gibon, Y., Bolger, M., Arrivault, S., Usadel, B., et al. (2011). Identification of Enzyme Activity Quantitative Trait Loci in a Solanum lycopersicum x Solanum pennellii Introgression Line Population. Plant Physiol. 157, 998–1014. doi:10.1104/pp.111.181594.

Szafranek, B. M., and Synak, E. E. (2006). Cuticular waxes from potato (Solanum tuberosum) leaves. Phytochemistry 67, 80–90. doi:10.1016/j.phytochem.2005.10.012.

Team, R. D. C. (2015). R: A Language and Environment for Statistical Computing.

Tipple, B. J., Berke, M. a, Doman, C. E., Khachaturyan, S., and Ehleringer, J. R. (2013). Leaf-wax n-alkanes record the plant-water environment at leaf flush. Proc. Natl. Acad. Sci. U. S. A. 110, 2659–64. doi:10.1073/pnas.1213875110.

Tipple, B. J., and Pagani, M. (2010). A 35Myr North American leaf-wax compound-specific carbon and hydrogen isotope record: Implications for C4 grasslands and hydrologic cycle dynamics. Earth Planet. Sci. Lett. 299, 250–262. doi:10.1016/j.epsl.2010.09.006.

Toubiana, D., Semel, Y., Tohge, T., Beleggia, R., Cattivelli, L., Rosental, L., et al. (2012). Metabolic Profiling of a Mapping Population Exposes New Insights in the Regulation of Seed Metabolism and Seed, Fruit, and Plant Relations. PLoS Genet. 8, 1–22. doi:10.1371/journal.pgen.1002612.

Warnock, S. J. (1991). Natural Habitats of Lycopersicon Species. HortScience 26, 1–6.

Yeats, T. H., Buda, G. J., Wang, Z., Chehanovsky, N., Moyle, L. C., Jetter, R., et al. (2012). The fruit cuticles of wild tomato species exhibit architectural and chemical diversity, providing a new model for studying the evolution of cuticle function. Plant J. 69, 655–66. doi: 10.1111/j.1365-313X.2011.04820.x.

